# Proteomic profiling across early postnatal development of midbrain dopamine axons innervating the dorsal striatum

**DOI:** 10.64898/2026.09.17.752390

**Authors:** Meghan Masotti, Lambertus Klei, Akayla Lewin, Vasin Dumrongprechachan, Bernie Devlin, Matthew MacDonald, Yevgenia Kozorovitskiy

## Abstract

Precise wiring and tuning of dopamine axons in the brain is crucial for normal development of basal ganglia circuits. This process takes place both pre and postnatally, with early postnatal windows reflecting important periods of terminal maturation. Protein expression and precise localization are vital to this process, underlying changes in axon structure and function. Yet, technical hurdles have prohibited mapping protein expression and localization as dopamine axons mature to form functional release sites. Here we employ targeted proximity labeling in midbrain dopamine neurons and track isolated distal axonal proteins in the striatum across development, from the neonatal period through adolescence. Our approach enabled high-precision evaluation of the axonal proteome, supported by a combination of Cre-recombinase based restriction and spatial separation. We map co-varying subpopulations of axonal proteins differentially expressed across development. Microtubule-associated proteins decreased over time, revealing temporal dynamics of dopaminergic axon stabilization at the proteomic level. Conversely, proteins supporting coordinated action potential firing increased over time, likely contributing to the characteristic tonic firing of dopamine neurons emerging over development. Intersecting our findings with recent GWAS of psychiatric and neurological disorders provided functional confirmation of disease and disorder risk. This analysis highlighted early expression and localization of risk gene products, providing spatial-temporal points of convergence for disease-associated variants specifically to the initial stages of postnatal neurodevelopment of dopamine neurons. Together, these findings provide the essential grounding for understanding postnatal dopaminergic axon development and contextualize neurodevelopmental functions for numerous risk gene products.

## Introduction

Neuromodulatory inputs, including monoaminergic systems, are essential for neurodevelopment and for fine-tuning the activity and plasticity of neurons throughout cortical and subcortical regions. In particular, dopamine is essential for refining basal ganglia circuits and for the development of motor control, reward-based learning, and decision-making (1,2). Centrally released dopamine originates from midbrain dopaminergic (DA) centers. Dopamine axons are present throughout the brain at birth, including in the striatum, where DA modulation of excitatory inputs is critical for basal ganglia function. Despite abundant axonal innervation at birth, canonical tonic DA release is not observed in rodents until postnatal days/P5-11 (3,4). Further, dopamine axons cluster into striasomes during early postnatal periods but gradually disperse throughout the striatum later. Mature DA striatal arbors become remarkably morphologically complex, with dense and highly branched terminals comprising ∼2-5% of striatal volume (5,6). This morphology is thought to contribute to high metabolic demands of these neurons, leading to vulnerability in psychiatric and neurological conditions (7,8).

Despite decades of study of the dopamine system, little is known about the underlying proteomic changes driving these postnatal transformations in DA neurons. Precise control of protein expression, localization, and turnover is critical across maturation. Understanding the proteome of DA neurons is therefore important for understanding both typical neurodevelopment and numerous diseases that involve the DA system. Recent advances in proximity labeling now allow for proteomic profiling of cell types and compartments in complex tissue. Through exogenous expression of engineered enzymes, nearby proteins can be tagged with small molecules, then enriched and quantified using mass spectrometry. One such enzyme, the engineered peroxidase APEX2, was designed to provide spatially specific biotinylation within seconds of induction, an advantage compared to labeling enzymes that operate on a timescale of minutes to hours (9–12). Furthermore, enzyme expression can be genetically targeted to cell types, via the Cre lox system, and subcellularly enriched within sub-compartments (13–15).

Here, we employed APEX-mediated proximity labeling to understand the dynamics of the developing dopamine axonal proteome, from the early postnatal period to adolescence. We restricted APEX expression to midbrain DA neurons and selected for axonal proteins by microdissecting striatum, quantifying proteins across development of these distal axons and clustering them based on developmental trajectories.

## Methods

Details for a subset of procedures, including mouse strains and husbandry, immunofluorescence staining, western blot, imputation, enrichment and developmental analyses, and data visualization are in **Supplementary Methods**.

### Proteomic tissue isolation

Sample collection follows Dumrongprechachan *et al., 2022* (13,14). Briefly, mice were deeply anesthetized with isoflurane. Brains from P6, P11, P20, and P50 animals were dissected, mounted on a Leica vibratome, and sectioned at 300 μm. Acutely prepared slices were incubated in artificial cerebrospinal fluid (ACSF: 127 mM NaCl, 25 mM NaHCO_3_, 1.25 mM NaH_2_PO_4_ monobasic monohydrate, 25 mM glucose, 2.5 mM KCl, 1 mM MgCl_2_, and 2 mM CaCl_2_, bubbled with carbogen (95% O_2_, 5%CO_2_). Slices were incubated in 500 μM biotin phenol (Iris Biotech, LS-3500.5000, lot: 1616162) diluted in ACSF for 1 hour (RT), bubbled with carbogen. Labeling was induced by 0.03% hydrogen peroxide for 2 minutes prior to quenching in 20 mM sodium ascorbate (Sigma Aldrich, A4043-100G, lot: SLBR2743V), both in ACSF. Dorsal striatal tissue punches (d=2 mm) were collected and flash-frozen prior to long term storage at -80°C until use.

### Sample preparation for mass spectrometry analysis

Tissue punches were thawed on ice and resuspended in 300 μL of mass spectrometry grade 1% SDS lysis buffer with HALT protease inhibitors and lysed with probe sonication for 20-30 seconds. Lysates were cleared by centrifugation at 12,000 RPM for 10 minutes (4°C) and transferred to a fresh low-binding tube (ThermoFisher Scientific, Waltham, MA). Samples were randomly assigned for subsequent steps, to mitigate order-based effects. Protein concentration was estimated using the BCA assay, with 500 μg of sample lysate in a final volume of 250 μL. Lysates were reduced with 20 μL 200 mM Dithiothreitol (DTT, Milipore Sigma, 10197777001, lot:77280321) for 30 minutes with shaking at 400 RPM and alkylated for 45 minutes with 60 μL 200 mM iodoacetamide (IAA, Sigma Aldrich, I1149, lot: SLC6164) in the dark with shaking.

150 μL of streptavidin beads (ThermoFisher Scientific, 88817, lot: ZJ402214) were brought to RT and washed with 100 mM TEAB (ThermoFisher Scientific, T7408, lot: 102485159). Reduced and alkylated lysates were added to beads and shaken at 1,200 RPM for 1.5 hours (RT). Beads were washed with 1 mL each: 2x with no-SDS lysis buffer (125 mM TEAB, 75 mM NaCl), 1x with 1 M KCl, and 5x with 100 mM TEAB. 0.4 μg trypsin resuspended in 150 μL of 100 mM TEAB was added to the enrichment reaction, overnight at 37°C.

Tryptic peptides were collected. Beads were washed with 40 μL of 100 mM TEAB. Wash supernatant and tryptic peptides were combined, then dehydrated and frozen at -80°C. Dehydrated peptides were thawed on ice and resuspended in 24 μL 100 mM TEAB, and bath-sonicated at RT for 15 minutes. For each sample, 4 μL of peptides was reserved prior to labeling, then combined to generate two pooled controls.

TMTPro reagents (ThermoFisher Scientific, A44521 lot: ZD385222) were thawed and resuspended in 18 μL of acetonitrile (ThermoFisher Scientific, A996-1, lot: 242509). TMT reagents were shaken for 5 minutes (RT). TMTPro reagent was added to samples (6 μL/sample). For two pooled controls, 12 μL of TMT reagent was added. TMT channel assignments were randomized. Labeling reaction was 1 hour-long, with shaking (RT). Reaction was quenched with 3 μL of 5% hydroxylamine/sample (ThermoFisher, 90115, lot: ZC389604) for 15 minutes, with shaking. For the two pooled controls, reactions were quenched with 6 μL of 5% hydroxylamine. Samples were combined based on plex assignments. Pooled controls were split between plexes. Plexes were frozen, then peptides were dried and stored at -80°C.

### Mass spectrometry data acquisition and raw data processing

Mass spectrometry data acquisition and processing was adapted from Dumrongprechachan *et al*., 2022. Briefly, TMT-labeled peptides were thawed and resuspended in 2% acetonitrile/0.1% formic acid, loaded onto a PepMap RSLC C18 column (Thermo Scientific, Waltham, MA), and eluted over 8 ACN gradients for high-pH reversed-phase fractionation.

For analysis, each fraction eluate was electrosprayed at 2,000 V into a Thermo Scientific Orbitrap Eclipse mass spectrometer (Thermo Scientific, Waltham, MA). MS1 spectra were acquired at 120,000 resolving power, and the Ion Trap with collision-induced dissociation (CID) of 35% in centroid mode was used to obtain MS2 spectra. To select ions for synchronous precursor selection in MS3, real-time max search parameters were set as follows: maxtime=34 s, max missed cleavages=1, Xcorr=1, dCn=0.1, ppm=5. The Orbitrap with HCD set to 60% was used to acquire MS3 spectra, with an isolation window=0.7 m/z, resolving power of 60,000, and max injection time of 400 ms.

Raw MS files were processed in Proteome Discoverer v2.4 (Thermo Scientific, Waltham. MA). Spectra were searched against the *Mus musculus* Uniprot/SwissProt database, using SEQUEST search engine. The following settings were used in the search: enzyme=trypsin, max. missed cleavage=4, min. peptide length=6, precursor tolerance=10 ppm. Included static modifications: carbamidomethyl (C,+57.021 Da), and TMT labeling (N-term and K,+304.207 Da for TMTpro16). Dynamic modifications included: (M,+15.995 Da), phosphorylation (S, T, Y,+79.966 Da), acetylation (N-term,+42.011 Da), Met-loss (N-term, –131.040 Da), and Met-loss + Acetyl (N-term, –89.030 Da). PSMs were filtered by the Percolator node (max Delta Cn=0.05, target FDR (strict)=0.01, and target FDR (relaxed)=0.05). Proteins were identified with a minimum of 1 unique peptide and protein-level combined q values<0.05, unless peptides were absent in >50% of Cre+ samples. Reporter ion quantification was based on corrected signal-to-noise values at these settings: integration tolerance=20 ppm, method=most confident centroid, co-isolation threshold=70, and SPS mass matches=65. PSMs and peptide level abundances from Proteome Discoverer were exported for analysis.

### Data processing

Data were exported from Proteome Discoverer and filtered at the peptide level. This yielded a total of 4,913 unique peptides. We removed 726 and 651 peptides not present in the pooled control samples, or in under half of the samples for each developmental stage. Peptides that were quantified in Cre+ but not in Cre-samples, were retained (2,552). For the 984 peptides quantified in both Cre+ and Cre-samples, 942 with a Log2FC >2 enrichment over Cre-background were retained. In addition, 23 peptides that were not quantified in any P6 samples and were the only peptides mapping to a given protein were excluded. These steps retained 3,741 peptides. Data were imputed using scVAEIT, as we have done before for transcriptomic and proteomic data (16) (see **Supplementary Methods**).

## Results

### APEX expression in midbrain dopaminergic neuron somata and projections across development

We used our previously generated floxed APEX reporter line (**Figure 1A**), where cytosolically directed, Cre-dependent APEX had been inserted into the permissive *Gt(ROSA)26* locus (17). EGFP is in frame with APEX, separated by a P2A linker, to facilitate characterization of APEX expression (14). To restrict APEX expression to dopaminergic neurons, we crossed the APEX reporter line to the DAT^iCre^ mouse line, where Cre recombinase expression is controlled under the endogenous DAT promoter (18). To confirm APEX expression and activity in DA neurons we induced APEX labeling in fixed tissues at P20 (**Methods**). Sections were co-stained for biotinylated proteins using streptavidin, and antibodies for eGFP and tyrosine hydroxylase (TH), the canonical marker for DA neurons and the enzyme that catalyzes the rate-limiting step in dopamine synthesis, respectively (**Figure 1B**). Nearly all TH-positive cells (>99%) expressed eGFP and were co-labeled by streptavidin (**Supplementary Figure 1**). To confirm APEX expression in striatal axonal processes, we induced APEX labeling in fixed tissue and co-stained with streptavidin and an antibody for DARPP32, a marker for spiny projection neurons. We observed expected extra-somatic but no somatic striatal labeling by streptavidin (**Figure 1C**), confirming specificity of APEX expression and activity.

**Figure 1:**
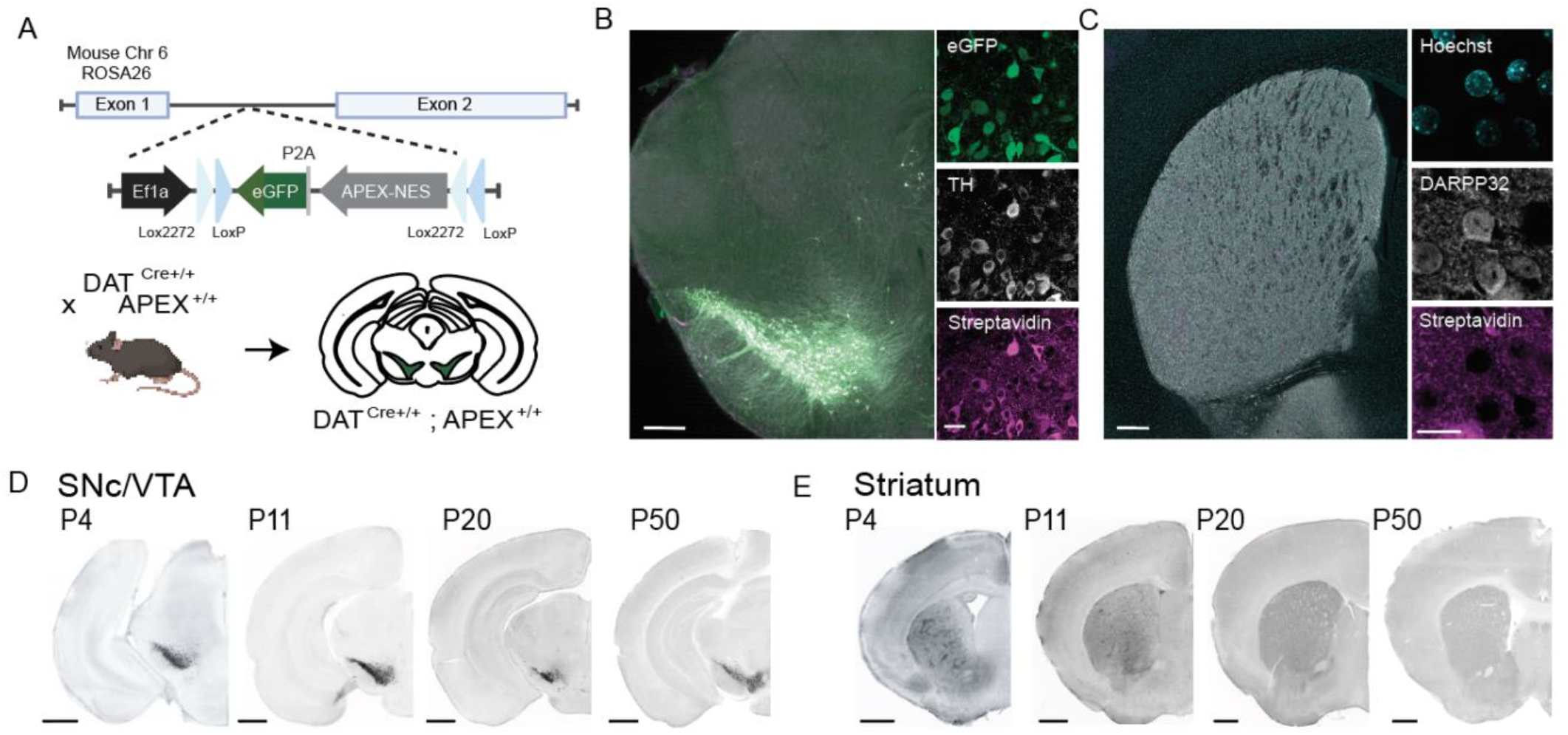
Cre-dependent expression of APEX in DA neurons across postnatal development. A) Schematic of genetically targeted APEX expression. *Top:* Cre-dependent APEX transgene design in the proteomic reporter mouse line, under the control of the EF1ɑ promoter. APEX is flanked by two loxP sites. GFP is in frame with APEX, separated by a P2A linker. *Bottom:* A cross to DAT^IRES*cre*^ expressing mouse line generates restricted APEX expression in DA neurons. B) Cre-dependent APEX expression in the VTA/SNc of a P20 DAT^IRES*cre* +/-^ ; APEX^+/-^ mouse. *Left*, image of immunolabeling for eGFP (green), tyrosine hydroxylase (grey), and streptavidin (magenta). Scale bar, 500 µm. *Right*, close-up confocal image of the SNc. Scale bar, 15 µm. C) Cre-dependent APEX expression in striatum of P20 DAT^IRES*cre* +/-^ ; APEX^+/-^ animal. *Left* image of Hoechst stain(cyan), immunolabeling for DARPP32 (grey), and streptavidin (magenta) Scale bar 500µm. *Right:* confocal close-up confocal image with additional 4x optical zoom of striatum. Scale bar 15µm. D) APEX expression in the SNc/VTA across periods of postnatal development. Immunolabeling for eGFP (greyscale). Scale bar, 100 µm. E) Same as D, but for the striatum.

We next sought to characterize APEX expression across early postnatal timepoints. We isolated tissue sections across P4, P11, P20, and P50, and immunostained for eGFP as a marker of APEX-positive regions. Throughout development we observed robust eGFP signal in the VTA/SNc. GFP immunolabeling in the striatum appeared spatially clustered during early development––likely concentrated in striosomes (6,19,20)—then became diffuse over time (**Figure 1C, D**).

### Selective enrichment of biotinylated proteins from dopamine axons innervating the dorsal striatum across development

We employed the APEX reporter line to selectively enrich proteins from dopamine axons projecting to the striatum, to identify and quantify them by bottom-up mass spectrometry. Four time points were selected based on developmental relevance: neonatal (∼P6), early postnatal (∼P11), preweanling (∼P20), and adolescent (∼P50). These timepoints encompass periods of DA neuron apoptosis, synaptic maturation, and pruning (21–25). We isolated dopaminergic axon proteins from acutely prepared striatal sections (Methods) (13–15). Striatum was micro-dissected for spatial isolation of axons. We confirmed the presence of biotinylated proteins across all time points via streptavidin labeling on western blot, using REVERT total protein stain as a loading control. Cre-negative controls were included to evaluate endogenous biotin labeling and bead background independent of APEX expression. Only Cre-positive samples showed labeling across a range of molecular weights (**Figure 2B)**.

**Figure 2:**
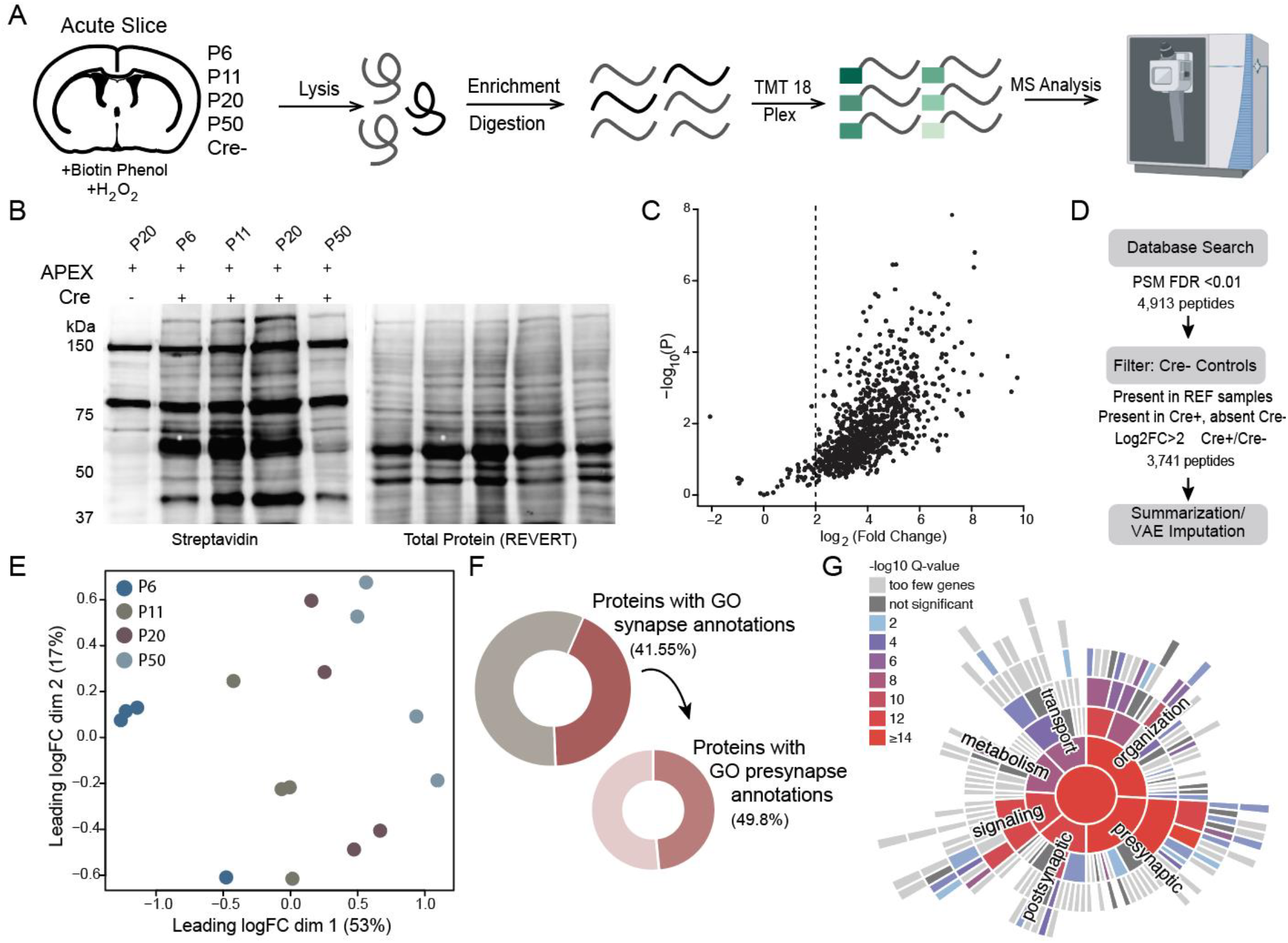
Proteomic profiling of DA axons projecting to the striatum. A) Schematic of sample collection for TMT multiplexed protein mass spectrometry analysis. B) Western blot analysis of biotinylated proteins in the striatum across development. Labeling was induced in 300 µm acutely prepared slices from P6, P11, P20, and P50 DAT^I-C*re* +/-^ ; APEX^+/-^ animals. P20 APEX^+/+^ ; DAT ^wt/wt^ tissue is included as no-label control. *Left:* streptavidin labeling. *Right:* Revert total protein stain. C) Volcano plot showing differential abundance of shared peptides between Cre-controls and experimental animals. X-axis represents Log2FC, y-axis represents -log10(p-values). Dotted line represents Log2FC of 2. D) Initial data processing and filtering. MS spectra were analyzed in Proteome Discoverer with a FDR <0.01 for peptide spectral matches. Rows were normalized to TMT abundance. Quantified unique proteins were filtered compared to Cre-controls, Missing values were VAE imputed. Peptide abundances were summarized into protein log2 abundances. E) Multidimensionality scaling (MDS) representation of protein level sample variance. Teal, P6; grey, P11; purple, P20; light blue, P50. F) *Left:* Proportion of quantified proteins with GO synapse annotations. *Right:* Proportion of quantified synaptic proteins that have presynaptic annotations. G) Sunburst representation of cellular component terms for synaptically annotated proteins by SynGO. Color-coded based on -log10 (Q value) for overrepresented terms within the data set by hypergeometric test.

For proteomics analysis, we isolated biotinylated proteins from dopamine axons from 4 biological replicates per time point (2 males, 2 females). For the P6 time point only, pooling from two sex-and age-matched replicates was needed to increase total protein. To control for endogenously biotinylated or nonspecifically enriched proteins we included four Cre-negative, APEX transgene-positive samples, with the same labeling protocol (**Figure 2A**). Protein lysates from tissue sections were enriched on streptavidin magnetic beads, then digested off-bead to generate tryptic peptides. Individual samples and controls for plex variability were barcoded with randomly assigned tandem mass tags (TMT) labels for sample identification and then pooled by plex. Pooled samples were high pH reverse-phase fractionated to reduce complexity and enhance peptide identification depth.

Peptide identification and quantification was conducted in Proteome Discoverer (2.4). For summarization and comparison to Cre^-^ samples, we first retained 2,552 peptides that were observed in 2 or more biological replicates for each time point and also not present in 3 or more Cre^-^ controls. As a second filter, for peptides that were present in 2 or more Cre^-^ controls, we retained 942 peptides that showed log2FC>2 in experimental samples (**Figure 2C, D**). In total, 3,741 peptides were included for further analysis. Plex effects were removed first, then additional covariates were determined using multidimensional scaling (MDS) (26) to understand underlying structural variance of biological replicates. For this analysis, we included only peptides observed across all biological replicates after removing plex effects. We summarized peptide abundance into log2 protein abundance and renormalized it to lane abundance, reflecting total intensity of peptides/sample. MDS analysis of the renormalized data showed the leading variance dimension mapped onto age and accounted for 53% of variance (**Figure 2E)**. Missing values were imputed using established pipelines we have developed (**Supplementary** Methods, (16,27)).

SynGO (v20231201) confirmed that 41.55% of these proteins have synaptic annotations (**Figure 2F, G)**. We do not expect the majority of axonal proteins to have synaptic annotations, as the samples included vast axonal arbors, with only a portion accounting for presynaptic terminals. Of the 41.55% with synaptic annotations, 49.8% were presynaptic. We further investigated the fraction of proteins with postsynaptic annotations, and noted that some proteins, such as G_ɑi2_—an important messenger in G protein-coupled receptor signaling cascades (28)––are annotated for both pre- and post-synapse. Additionally, many proteins with exclusively postsynaptic annotations in SynGO have emerging evidence for presynaptic localization (29–31). For example, myosin IIB heavy chain protein (MYH10) is vital for postsynaptic cytoskeleton organization but has also been shown to have a role in presynaptic vesicle release (32). We hypothesize that multiple other proteins captured in our axonal enrichment have both pre- and post-synaptic function, highlighting the need for continued proximity labeling and advanced microscopy studies to refine our understanding of presynaptic proteomes across different circuits (14,15,33–36).

### Axonal proteins from dopamine neurons projecting to the striatum cluster based on developmental trajectory

We next examined how proteins change over development, from neonate to adolescence. Using the Flexmix R library (37), we fit age-based quadratic models to model protein trajectories (**Figure 3C**). To determine protein assignments to clusters, we implemented a posterior probability cut off of >0.8. All 1,197 proteins were assigned to a cluster, and average protein abundance within a trajectory over time was plotted (**Figure 3C, Supplemental Figure 3**). From this we determined that 279 proteins increased in abundance (grey), 575 remained at stable expression levels (light grey), while 343 decreased (dark grey).

**Figure 3:**
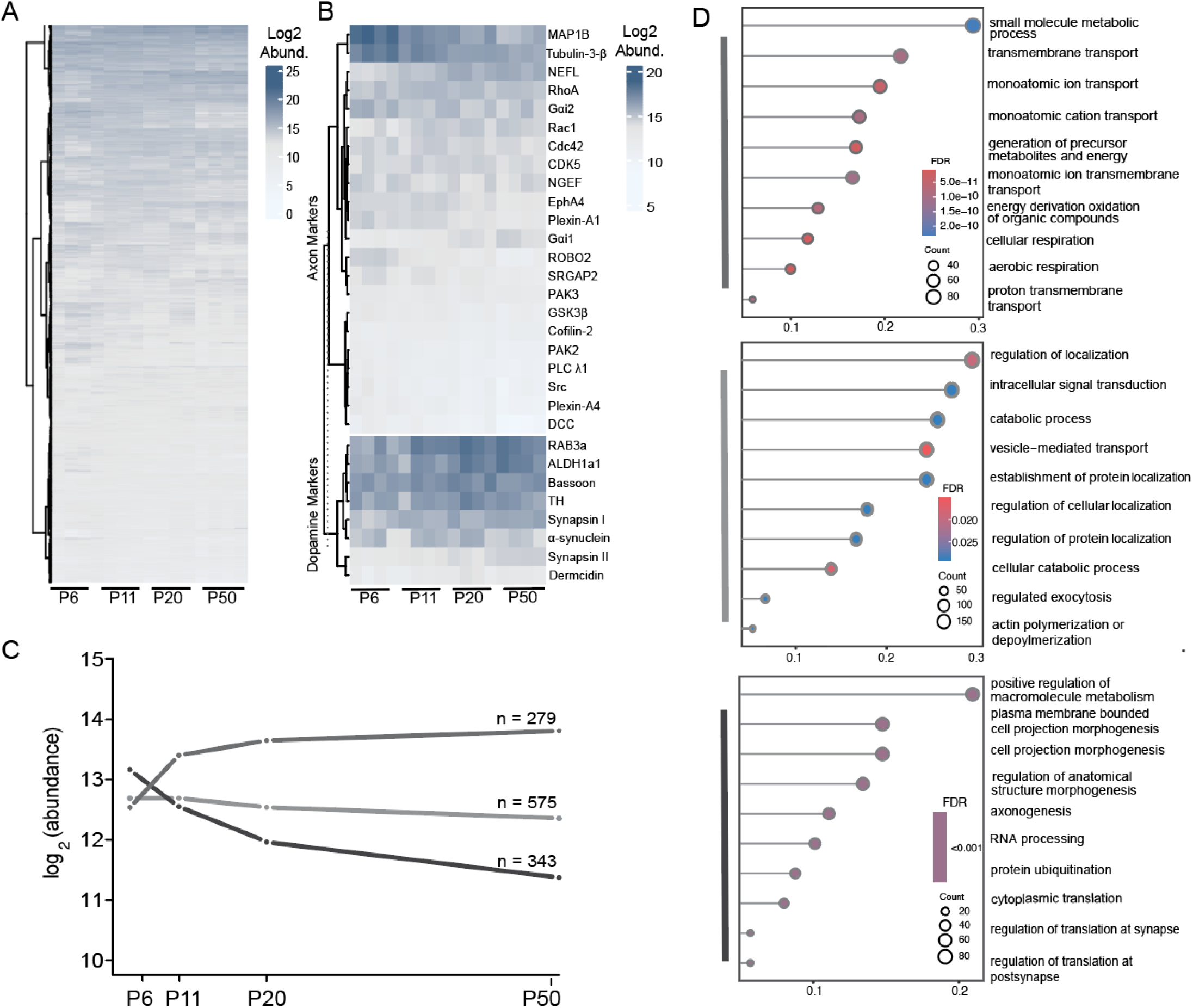
Temporal trajectories of protein abundance in DA axons. A) Heatmap of all quantified proteins across four time periods of postnatal development. Columns are ordered by developmental stage, rows are hierarchically clustered based on Euclidean distance with Ward D2 correction. Color map reflects Log2 normalized protein abundance. B) Same as A but for *top:* axon markers and *bottom:* dopamine neuron markers. C) Mean abundance for each group of clustered proteins, connected by lines between two developmentally sequential means. N reflects the number of proteins per group. Individual protein trajectories are plotted in Supplementary Figure 3. D) GO enrichment analysis for each trajectory cluster. All ontology annotations—cellular component, biological process, and molecular function—were included. Circles are ranked by GeneRatio, reflecting the fraction of genes found in this data set, compared to total genes in the GO term. Circle size and color reflect the number of genes and FDR (corrected p value) respectively. *Top:* Proteins with increasing trajectories. *Middle:* Proteins with stable trajectories. *Bottom:* Proteins with decreasing trajectories. The complete list of significant enrichment terms is in Supplemental Table 3.

To better understand what processes are regulated by these three trajectory groups, we performed GO analysis with R library clusterProfiler (38) (**Figure 3D, Supplementary Table 2**). For decreasing in abundance proteins, we noted enrichment for translation-associated terms, such as RNA processing, aligning with known developmental trajectories for local protein synthesis and synaptic plasticity (25,39,40). Machinery associated with vesicle release largely stayed constant, consistent with the literature showing dopamine release by birth or even prior, well before complete wiring of the basal ganglia circuits and the onset of phasic and tonic activity patterns (41). Interestingly, proteins corresponding to membrane transport and metabolism increased in abundance over time, potentially reflecting growing metabolic activity of dopamine neurons (7,42). Together, this suggests that while dopamine axons are present and functional immediately after birth, tonic firing patterns seen in adulthood are mediated by upregulation of ion channels and ion exchangers, rather than vesicle release and docking proteins. We also investigated the KEGG pathway terms enriched for each temporal cluster (**Supplemental Figure 3**) and noted enrichment for gene products associated with neurodegenerative diseases, as well as shared pathways for neurodegeneration in each temporal cluster, highlighting the vulnerability of dopamine neurons in neurological diseases.

### Overrepresented terms align with expected developmental trajectories

To further examine developmental protein groups identified by term enrichment analysis, we selected a high-confidence representative GO term from each trajectory cluster: transmembrane transport (GO:0055085) for increasing, vesicle-mediated transport (GO:0016192) for stable, and axonogenesis (GO:0007409) for the decreasing cluster, for additional unsupervised analysis. We isolated all DA axon-expressed proteins within those GO terms, and visualized entire group dynamics via heatmap (**Figure 4A**). We confirmed that GO group dynamics aligned well with the cluster they were overrepresented in. A small fraction of each GO group displayed differential changes in expression antagonistic to the overall group GO dynamic, highlighting the need to interrogate proteins within networks to understand their cellular functions.

**Figure 4:**
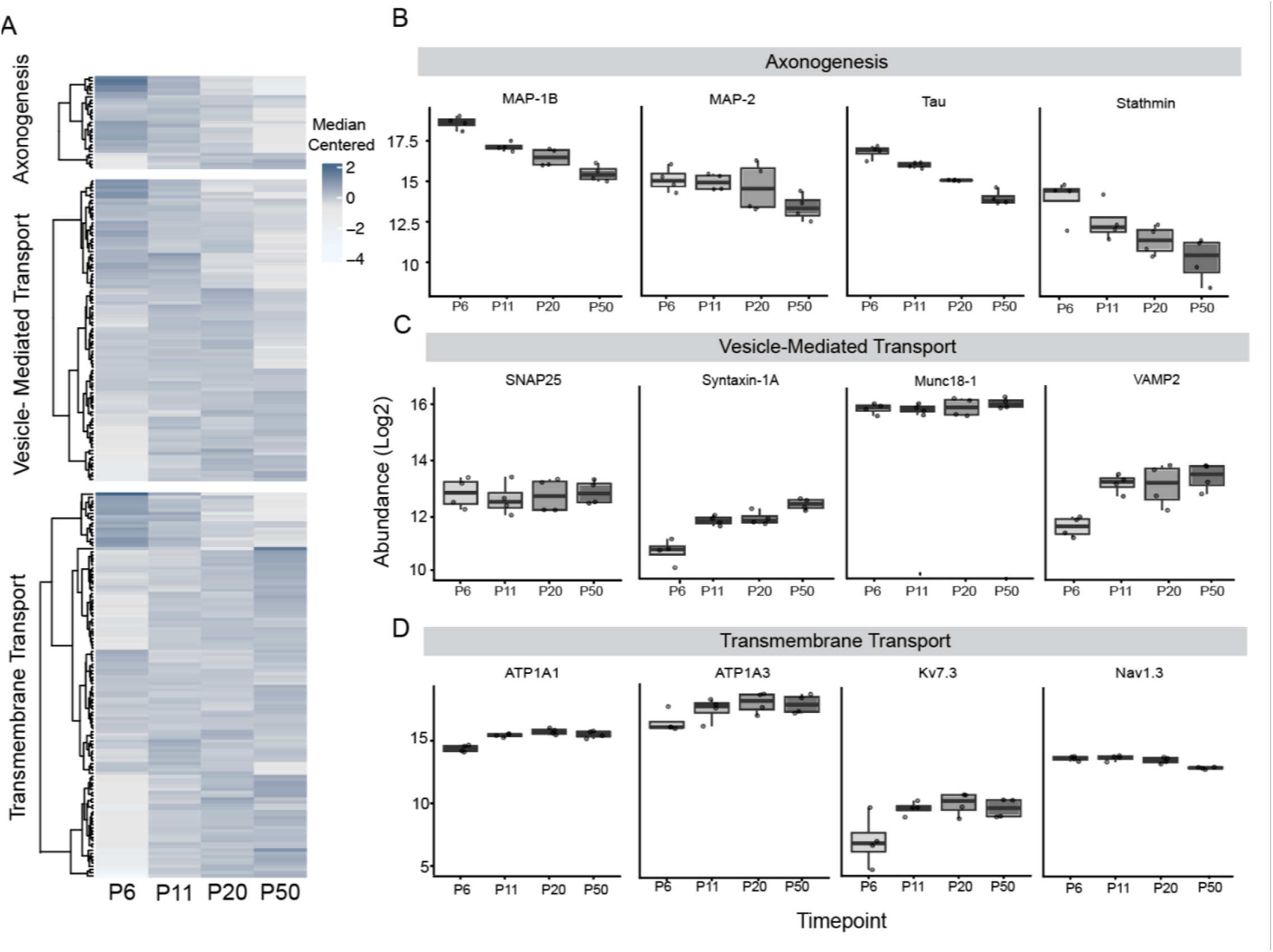
Over-represented terms align with axon development trajectories. A) Heat map of selected over-represented terms. Columns reflect the average protein abundance at each age, ordered by developmental stage. Rows are hierarchically clustered based on smallest Euclidean distance with Ward D2 correction. Data are median centered, and value is reflected in color map. B) Box plots of abundance for selected proteins from the GO term Transmembrane Transport (GO:0055085). Each point represents Log2 protein abundance from one biological replicate. C) Same as B, but for the Vesicle-Mediated Transport term (GO:0016192). D) Same as B, but for the Axonogenesis term (GO:0007409).

For mapping temporal dynamics of interacting protein sets, we isolated co-expressed or known to interact proteins within each GO group. Among those increasing expression, we noted several important ion channels (Na_v_1.3, K_v_7.3) and sodium potassium pump subunits (ATP1a1, and ATP1a3). All except Na_v_1.3 increased expression (**Figure 4D)**, suggesting that tools for ion movement follow the overall trajectory of dopamine axon maturation. Similarly, we isolated proteins with consistent expression over time. We chose to investigate proteins within the SNARE complex and their binding partners. While SNAP25, a crucial SNARE complex protein, and STXBP1, Syntaxin-binding protein 1 (Munc18-1), stably express across development, two components essential for vesicle release (STXA1 and VMP2) slightly increase as DA axons mature (**Figure 4C)**. Finally, we also examined microtubule-associated proteins. We hypothesized that several may decrease as axons mature (**Figure 4B)**, as synaptic plasticity decreases with age, although MAPT has been found to increase in expression in cultured neurons over time (43). Indeed, MAP1b, MAP2, and STM1 all decreased over time, but we also observed that MAPT (tau) decreased in expression.

### Proteins encoded by risk genes for psychiatric and neurological conditions in DA neurons

Disrupted dopamine signaling, including specifically dysregulation of dopaminergic axons, have been implicated in several neuropsychiatric and other disorders. We sought to characterize which associated genes translate into proteins during postnatal development, and whether they change. A list of risk genes for several conditions was generated using GWAS data from the NHGRI-EBI Catalog of Human Genome-Wide Association Studies (44), as well as a list of autism (ASD)-associated genes from the Simons Foundation for Autism Research (SFARI) (45). Other conditions we included were bipolar disorder (BD), schizophrenia, major depressive disorder (MDD), epilepsy, and Parkinson’s disease (PD). We then filtered for proteins encoded by associated genes quantified in our data, plotting the average abundance over development (**Figure 5A**). We quantified 211 axonal proteins associated with genetic association for ASD, 13 for epilepsy, 82 for BD, 120 for MDD, 150 for schizophrenia, and 38 for PD.

**Figure 5:**
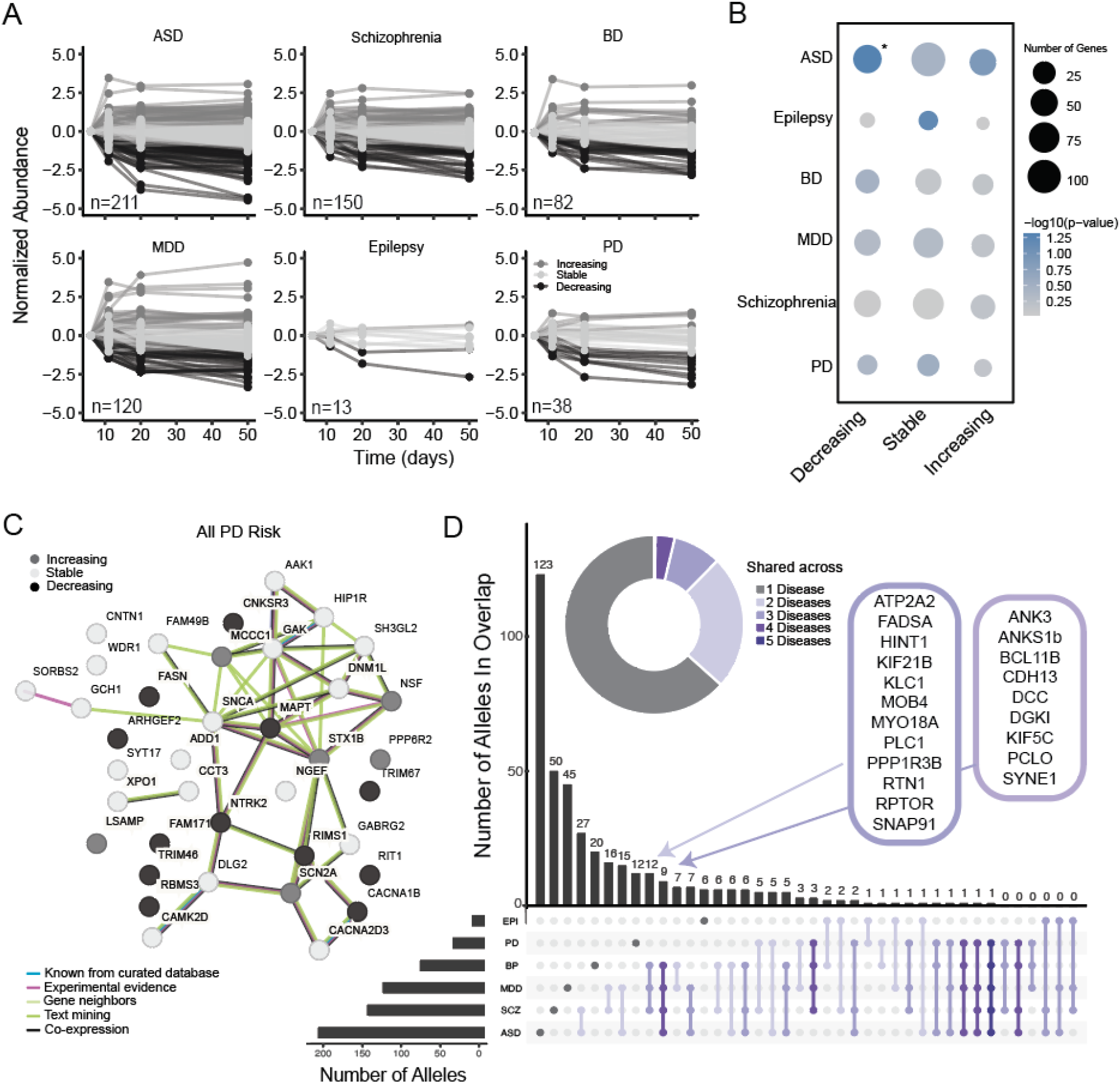
Temporal patterns of disease risk allele expression in DA neurons. A) Proteins were filtered for known risk alleles for Autism Spectrum Disorder (ASD), Bipolar Disorder (BP), Epilepsy, Major Depressive Disorder (MDD), Schizophrenia, and Parkinson’s Disease (PD). Data are color coded for trajectory clusters (grey, increasing; light grey, stable; dark grey, decreasing). B) Over-representation analysis for disease risk alleles within temporal clusters. Circle size indicates number of risk alleles found within each cluster, and color represents -log10(pvalue). C) String network analysis generated by String-DB for all PD risk alleles tracked across DA axon development. Circle color represents clusters in either increasing (grey), stable (light grey), or decreasing trajectories (dark grey). Line thickness represents the strength of interaction, and color reflects the evidence for interaction (red, gene fusion, green= neighborhood evidence, blue= co-occurrence evidence, purple = experimental evidence, black=co-expression evidence, and cyan= external database evidence). D) Upset plot showing shared risk alleles across diseases. Total identified risk genes in DA axons *(left*). Group comparison key (*bottom)*, with the number of shared genes identified *(graph bars)*. The numbers above each graph bar reflects the total number of shared genes between groups being compared. *Inset left:* donut chart depicting the number of risk alleles shared between multiple disease states. *Inset middle:* Shared risk alleles between Schizophrenia, MDD, and BP. *Inset right:* Shared risk alleles between Schizophrenia, MDD, BP, and ASD.

We then used R package clusterProfiler (38) to perform overrepresentation analysis to determine if trajectory groups were enriched for conditions (**Figure 5B**). Proteins encoded by ASD genes were overrepresented among those decreasing in expression over development (hypergeometric test, p = 0.048). Proteins encoded by risk alleles for PD and epilepsy did not reach statistical significance for overrepresentation in the stable trajectory (PD, p = 0.294; epilepsy, p = 0.063). Still, due to the large body of work highlighting SNc dopamine axon vulnerability in PD, we further examined whether the quantified proteins encoded by these genes were known interactors. We used stringDB (46) to generate a string diagram of all PD genes coding for proteins quantified in this data set (**Figure 5C**), and noted a central hub of interacting proteins primarily centered around MAPT (tau) and ɑ-synuclein.

We quantified several proteins encoded by genes shared across disorders. Notably, one protein—RIMS1, an important regulator of synaptic release—conferred liability for 5 conditions (ASD, BD, schizophrenia, MDD, and PD). We also identified a set of 16 proteins that confer risk for development of four conditions (**Figure 5D, Supplemental Figure 4**). We focused on shared genes for mood disorders (MDD, schizophrenia, BD), and noted 12 proteins from this group, including RPTOR, the regulatory subunit of mTORC1, which has been shown to modulate dopamine release and axonal morphology (47) (**Figure 5D, Supplemental Figure 4**). Overall, these data provide further evidence for dopamine system neurodevelopment as a point of convergence for genetic liability across both psychiatric and neurological conditions.

## Discussion

In this study, we expressed APEX across multiple developmental time points in midbrain dopamine neurons and implemented mass spectrometry-based bottom-up proteomics to identify axonal proteins via microdissection of the striatum in acutely prepared sections. We selected for axons as they are the source of dopamine release and comprise a large fraction of total neuronal volume, estimated to account for >80% of total protein absolute abundance (15). The Cre-dependent APEX reporter line facilitated access to early postnatal timepoints—a significant advantage over viral strategies that require weeks of incubation to robustly express proximity labeling enzymes. The ability of APEX to label rapidly, within seconds to minutes, facilitated capturing “snapshots” of cell state, without temporal blurring of protein expression.

We quantified 1,197 proteins across early postnatal development into adolescence, including several dopamine axon markers, including TH, AlLDH1a1, and α-synuclein (15). Among the quantified proteins, we determined that 41.55% had synaptic annotations. Proteins that change over development were associated with axon maturation and synapse stabilization. Known remodelers of axonal structure, such as the MAP protein family, decrease with age, consistent with decreasing axonal remodeling as terminals stabilize functional release sites (48–50). Conversely, action potential firing machinery, like sodium channels, increases over development. In parallel, sodium-potassium pump subunits, ATP1a1 and ATP1a3 (51,52), reach their peak expression around periods of rapid dopaminergic synapse maturation (P11-P15), corresponding to the advent of phasic, pulsatile dopamine release characteristic of these neurons. Mitochondrial proteins increased alongside sodium-potassium pump levels, likely to support increased firing. Proteins with stable expression over this postnatal window include those required for vesicle release, reflecting the ability of dopamine neurons to synthesize and release neurotransmitters at birth (3,41,53,54).

Dopamine neurons have long been implicated in in psychiatric and neurological disorders (55,56). Subsets of SNc dopamine neurons are the most vulnerable cell population in PD, and their loss relatively early in disease progression leads to deficits in motor control. Many proteins encoded by PD risk alleles fell within the stably expressing cluster, consistent with the emerging narratives on early vulnerability of dopamine neurons before disease onset (56–60). Dopamine dysregulation in the striatum has also been implicated in ASD. We quantified 211 proteins encoded by ASD genes within dopamine neuron axons alone (∼17.5% of all SFARI risk genes), highlighting the importance of axonal proteins in neurodivergent development. ASD gene products were overrepresented in the decreasing trajectory cluster, aligned with prior studies showing early brain-wide changes in neurodevelopmental conditions with bulk proteomics approaches (61,62). The ability of genetically restricted proximity labeling to target axonal subcompartments with high precision revealed temporal trajectories for numerous ASD gene products across diverse neuronal groups, including deep-layer cortico-striatal pyramidal cells (14) and dopamine neurons, as we find here. The distribution of load across both sides of the synapse is an exciting emerging area in understanding ASD risk loading across neural systems.

Several considerations and limitations merit acknowledgement. First, APEX expression was targeted throughout the neuron (14,15,63) and not specifically restricted to presynaptic terminals. Thus, we quantify the developmental trajectories of many structural axonal proteins, in addition to presynaptic machinery. Second, we cannot rule out a minor contribution of labeling of dendritic proteins via cell lysis events during acute slice isolation, which could free APEX to diffuse and label nearby compartments. Similarly, such lysis events and hydrogen peroxide treatment may trigger oxidative stress responses that could confound protein quantification, although the speed of labeling minimizes this concern. Despite these limitations, APEX-based approaches provide an important complement to other proximity labeling methods that biotinylate proteins *in vivo*. APEX supports quantification of precise and highly restricted snapshots of protein state.

Overall, this study was the first, to our knowledge, to isolate the early postnatal axonal proteome from dopamine neurons projecting the striatum across development. While individual proteins and neuronal properties have been extensively investigated in this population across development, and more generally, multiple genetically-targeted studies of specific neuronal proteomes have been carried out in adult mice, this work is the first to combine proteomics with dopamine axon development across key maturation ages. This begins to open the window into essential mechanisms that move in concert with respect to the proteomic abundance, along with genetic risk for varied brain disorders mapping onto these protein networks.

## Acknowledgements

This work was supported by Northwestern High Throughput Analysis Core. (RRID: SCR_017879). We are grateful to Lindsey Butler for mouse colony management. Some schematics were created with https://biorender.com/.

## Funding

This work was supported by the NSF CAREER Award 1846234, NIMH R56MH113923, NINDS R01NS107539, NIMH R01MH117111, the Beckman Young Investigator Award, Searle Scholar Award, Rita Allen Foundation Scholar Award, and Sloan Research Fellowship (all YK), and NIMH R01MH118497 (MLM). Research reported in this publication was supported by the Chemistry of Life Processes training grant, NIGMS T32GM105538 (MM).

## Author Contributions

**MMas-**Sample collection, preparation, experimental design, data analysis, manuscript writing. **LK-**Data analysis statistical oversight, manuscript edits. **AL-**Sample prep, data acquisition. **VD-**Sample collection, experimental planning. **BD-**Data processing and statistical oversight, manuscript editing. **MMac-**Data acquisition analysis oversight, manuscript edits. **YK-**Experimental design, analysis oversight, manuscript writing and editing.

## Conflict of Interest

The authors declare no conflict of interest.

## Notes

### Competing Interest Statement

The authors have declared no competing interest.

