## Supplementary Methods and Figures for "Proteomic profiling across early postnatal development of midbrain dopamine axons innervating the dorsal striatum"

### Supplemental Methods

#### Mouse strain and husbandry

All mice were handled according to IACUC approved protocols at Northwestern University. Mice were kept in humidity- and temperature-controlled, 12-hour light/dark cycle rooms, with standard enrichment and feeding protocols. Neonatal (P5-P6), early post-natal (P11-P12), preweanling (P19-P20), and adult mice (P48-P50) were used in this study. DAT$^{iCre}$ mice (Jax #006660, B6.SJL-Slc6a3tm1.1(cre)Bkmn/J, (18) and Cre-dependent APEX2 reporter mice (14) were used in this study. DAT$^{iCre+/+}$ were crossed with APEX2$^{+/+}$ mice to generate DAT$^{iCre+/-}$ ; APEX2$^{+/-}$ experimental animals. APEX2$^{+/+}$ mice were used as controls. Genotyping primers (DAT$^{iCre}$ - common F: TCGCTGTTGGTGTAAAGTGG DAT$^{iCre}$ mutant R: CCAAAAGACGGCAATATGGT, DAT$^{iCre}$ WT R: GGACAGGGACATGGTTGACT. APEX2- APEX F (R26F1(AAG)): CCAAAGTCGCTCTGAGTTGTTATCAGTAAG, APEX2 R (R26R2): TCTGTGGGAAGTCTTGTCCCTCC). Pups were randomly assigned to conditions.

#### Immunofluorescence staining

P11, P20, and P50 mice were deeply anesthetized on isoflurane prior to fix perfusion with 15-20 mL of 4% paraformaldehyde (PFA) in 0.1M phosphate buffered saline (PBS). P4 mice were deeply cryoanesthetized instead and not perfused. All brains were dissected and post-fixed in 4% PFA in PBS for 24-48 hours at 4°C. Brains were stored in PBS at 4°C until sectioning in PBS at 60µm thickness using a vibratome (VT1000s, Leica, Germany). P4 brains were agarose embedded in 1% low melting point agarose (Sigma Aldrich, lot: SLCD3511) in 0.1M PBS. Agarose was allowed to completely solidify around the brain prior to sectioning.

For biotin labeling in fixed tissues, sections were first incubated in 5mM biotin phenol (Iris Biotech, LS-3500.5000, lot: 1616162) shaking for 2 hours at RT, then labeling was induced with 0.5% hydrogen peroxide for 15 minutes. Sections were washed 3x 10 minutes in PBS, then permeabilized in 0.2% triton shaking at RT for 1-2 hours prior to incubation with streptavidin conjugated to Alexa Fluor 647 (Invitrogen S32257, lot: 2068269) overnight at room temperature. For antibody immunofluorescence, sections were permeabilized in 0.2% Triton-X diluted in PBS (PBS-T) and placed on a shaker at RT for 1-2 hours. They were then blocked in 5% bovine serum albumin (BSA) with 0.1% Triton-X in PBS shaking for 1 hour. Primary antibodies were diluted 1:500 in in 5% BSA with 0.1% Triton-X in PBS and incubated at 4°C overnight. Primary antibodies: Abcam chicken anti-GFP (AB13970, lot: 101873-40); Abcam rabbit anti-tyrosine hydroxylase (AB152, lot: 3595105); Cell Signaling rabbit anti-DARPP32: (19A3, lot: 2306S)**.** Sections were then washed 3 times for 10 minutes in PBS, with continuous shaking. Secondary antibodies were diluted 1:5,000 in PBS-T, and sections were incubated for 1-1.5 hours at RT with mild shaking. Secondary antibodies (all Invitrogen): goat anti-chicken conjugated to Alexa Fluor 488, (A11039, lot: 2566343); goat anti-rabbit conjugated to Alexa Fluor 594 (A11012, lot: 2616076); streptavidin conjugated to Alexa Fluor 647 (S32257, lot: 2068269). Sections were washed for 10 minutes 3 times in PBS with continuous shaking. Sections were mounted on Superfrost Plus slides (ThermoFisher Scientific, Waltham, MA), dried at room temperature in the dark, and then coverslipped in glycerol mounting media (3:1, glycerol:PBS) with 2 µg/mL Hoechst 33,342 (ThermoFisher Scientific; H1399, lot: 1611090). Slides were imaged using a 10x air objective using a VS120 slide scanning microscope (Olympus/Evident Scientific, Waltham, MA) or with a 60x oil objective using Floview Confocal (Olympus/Evident Scientific, Waltham, MA)

#### Western blot

Tissues were thawed on ice. SDS lysis buffer (1% SDS, 125 mM TEAB, 75 mM NaCl) was supplemented with HALT protease inhibitors (ThermoFisher Scientific, 1861282, lot: AC409694). 200 µL of lysis buffer was added per 20 mg of tissue. Tissues were lysed with probe sonication for 20-30 seconds. Lysates were cleared by centrifugation at 12,000 RPM for 10 minutes at 4°C. Cleared lysates were transferred to a fresh low-binding tube for storage and use (ThermoFisher Scientific, Waltham, MA).

Protein concentration was estimated using Bicinchoninic Acid (BCA) assay (ThermoFisher Scientific, Waltham, MA). Samples were diluted either 1:100 or 1:50 in a 96-well plate prior to quantification. Reagents were mixed per manufacturer specifications, for an incubation at 37°C for 30 minutes, or for 1-2 hours at RT. Fluorescence measurement was obtained at 562 nm using a SynNeo2 Plate reader (BioTek, Winooski, VT).

20-30 μg of protein was diluted in the lysis buffer and supplemented with 6x laemeli buffer (375mM Tris-HCl (pH 6.8), 9% SDS, 50% glycerol, 0.03% bromophenol blue) with 10% beta-mercaptoethanol. Protein was denatured at 95°C for 10 minutes and run on a 12% PAGE gel, at 90 V for 2 hours in standard SDS running buffer (1% SDS, 25 mM Tris base, and 192 mM glycine). Gels were transferred to nitrocellulose membrane (926–31090, LI-COR, NE, USA) in ice-cold transfer buffer (25 mM tris base and 192 mM glycine) for 1 hour at 75 V. Membranes were briefly washed in 1 M tris-buffered saline (TBS) and incubated overnight at 4°C in Streptavidin CW800 (LI-COR, 926-32230, lot: D10311-05), diluted 1:10,000 in TBS. Membranes were washed 3 times for 10 minutes in TBS. REVERT protein stain (LI-COR) was used to image total protein according to manufacturer instructions. Membranes were imaged on a LI-COR odyssey CLx scanner.

**Proteomic data processing: imputation**

After filtering, as described in the **Methods**, imputation was performed on the remaining 3,471 peptides using a probabilistic variational autoencoder model, scVAEIT, as has been done in proteomics and transcriptomics studies (16). ScVAEIT is a two-step process, where starting values are calculated using softImpute and information from the reference samples. This was followed by the imputation in scVAEIT, using 16 samples across the 4 developmental timepoints. Imputed peptide abundances were normalized to the average total sample abundance. Sample-specific weights were calculated by taking the average total abundance across samples and dividing this by the sample-specific total. Each peptide abundance was subsequently multiplied by the sample-specific weight. After removing 257 peptides which mapped to multiple proteins, the peptide-level data were aggregated into protein-level data for 1,197 proteins by summing peptide abundances for each protein. After aggregation, protein-level data were renormalized to the average total sample abundance. Sample specific weights were calculated as for peptides, where the average total abundance of each sample was divided by the sample-specific total.

**SynGO enrichment analysis**

To evaluate synaptic annotations, proteins that pass the Cre-negative filter described above were used for SynGO enrichment analysis (version 20231201). Entries corresponding to multiple isoforms of the same protein were collapsed into a single protein-level entry. Protein IDs were converted using the SynGO online converter tool. Graphs were downloaded and exported to Adobe Illustrator for figures.

**Developmental clustering**

The function flexmix, from the R package flexmix (37), was used to fit quadratic functions to $\log_2$ imputed protein-level data, adjusted for the effect of plex. We chose to use k=3 to model three main trajectories over developmental time: stable, increasing, and decreasing. Trajectory identity, assigned by flexmix, was based on a posterior probability exceeding 0.5.

**GO enrichment and disease risk gene overrepresentation analyses**

To test overrepresented terms in each developmental cluster, the R package clusterProfiler 4.0 (38) was used. Protein IDs were converted from UniProt/SwissProt to Gene symbols using BioMart (64). Each group (stable, increasing or decreasing) was analyzed using enrichGO function, against the background of all identified proteins. Ontology was set to all; p values were Benjamini-Hochberg corrected. Overrepresentation of KEGG pathways was also determined using clusterProfiler, with a q value $< 0.05$.

Disease risk gene overrepresentation (ORA) analysis was performed using GWAS data downloaded from the NHGRI-EBI Catalog of Human Genome-Wide Association Studies (access date 02-16-2026) (44) and SFARI ASD gene list (access date 3-15-2026) (45). Mouse symbols were converted to human symbols using the Uniprot idMapping tool. ClusterProfiler was used to determine the overlap between mapped genes in the GWAS data sets and protein-level data for each developmental group. The universe was set to all identified proteins. Hypergeometric tests were performed in clusterProfiler to determine if the overlaps were significant, with p value $<0.05$.

**Data Visualization**

Graphical representations were generated in R through either clusterProfiler (38) or ggplot2 (65), then exported as PDFs into Adobe Illustrator (San Jose, CA, ver. 2021) for further processing and arrangement into figures. Confocal and wide field images were exported into FIJI/ ImageJ (Bethesda, MD) for processing.

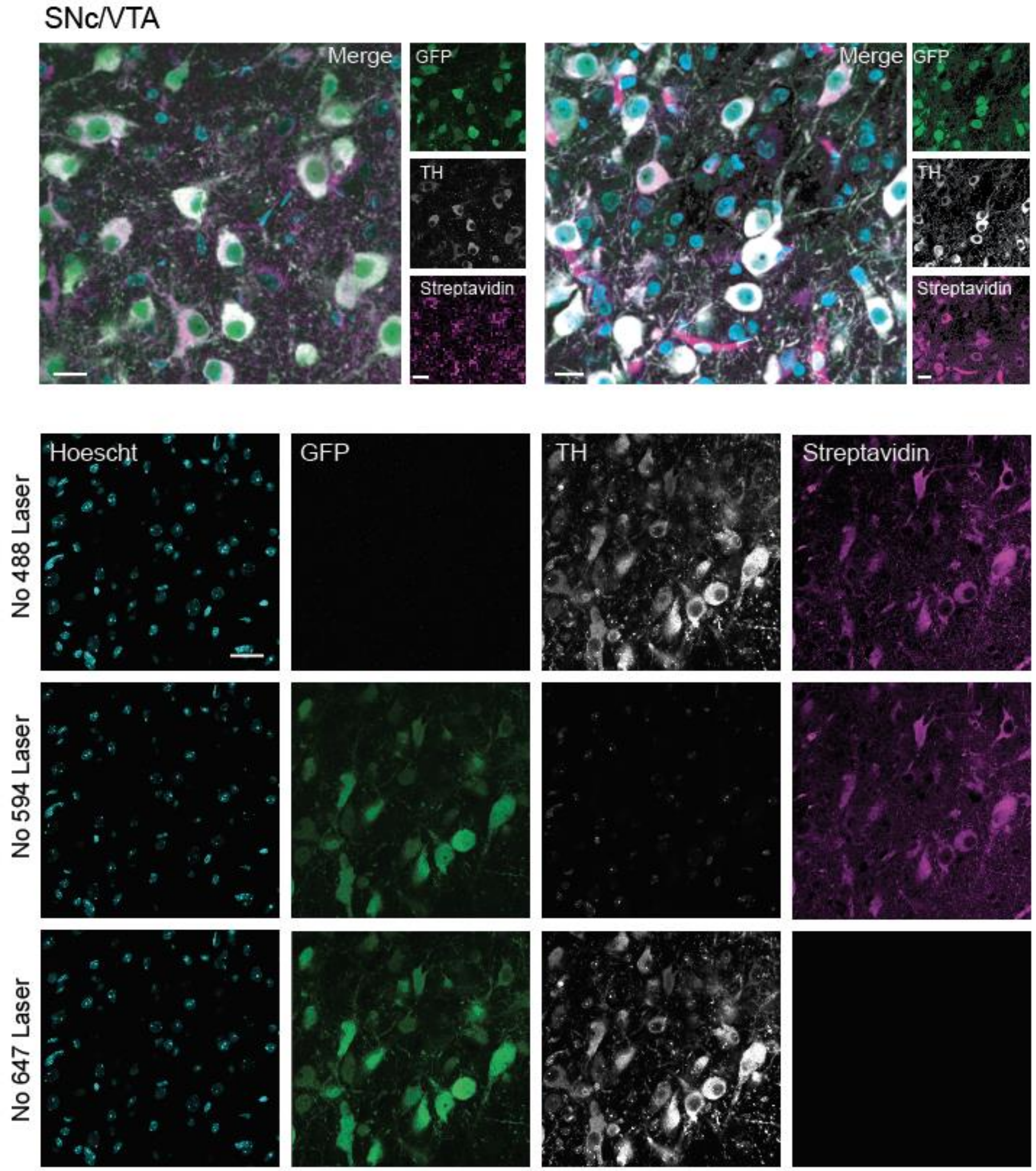


**Supplemental Figure 1: Sample confocal images of APEX biotinylation in DA neurons**

A) Representative images used for overlap quantification of TH+ cells in the SNc/VTA with GFP and Streptavidin. Merge image of Hoechst (imaged with a 405 nm laser), immunolabeled for GFP (imaged with a 488 nm laser), immunolabeled for TH (imaged with a 597 nm laser), and immunolabeled for Streptavidin (imaged with a 647 nm laser). Scale bars, 15 µm.

B) Sequential imaging of the same field of view within the SNc/VTA to determine the extent, if any, of spectral bleed between channels. All images are taken at the same magnification with the same laser power, except for indicated laser shut off, and post processed in the same way. Scale bars, 15µm.

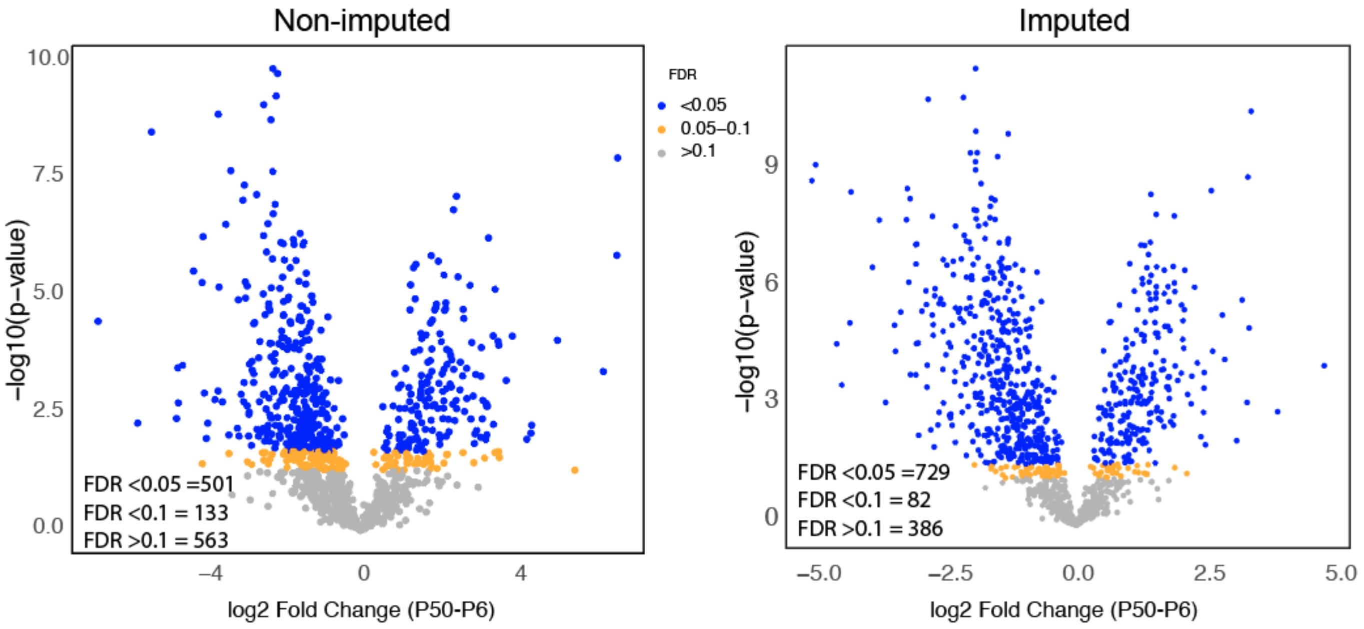


**Supplemental Figure 2: Comparative data pre- and post- imputation**

Comparison of fold change between P6 and P50 for non-imputed (*left),* vs imputed (*right)* data. Linear model-based p value, FDR, and fold-change calculation. Yellow dots represent proteins with q < 0.05, and blue dots represent proteins with q value <0.1.

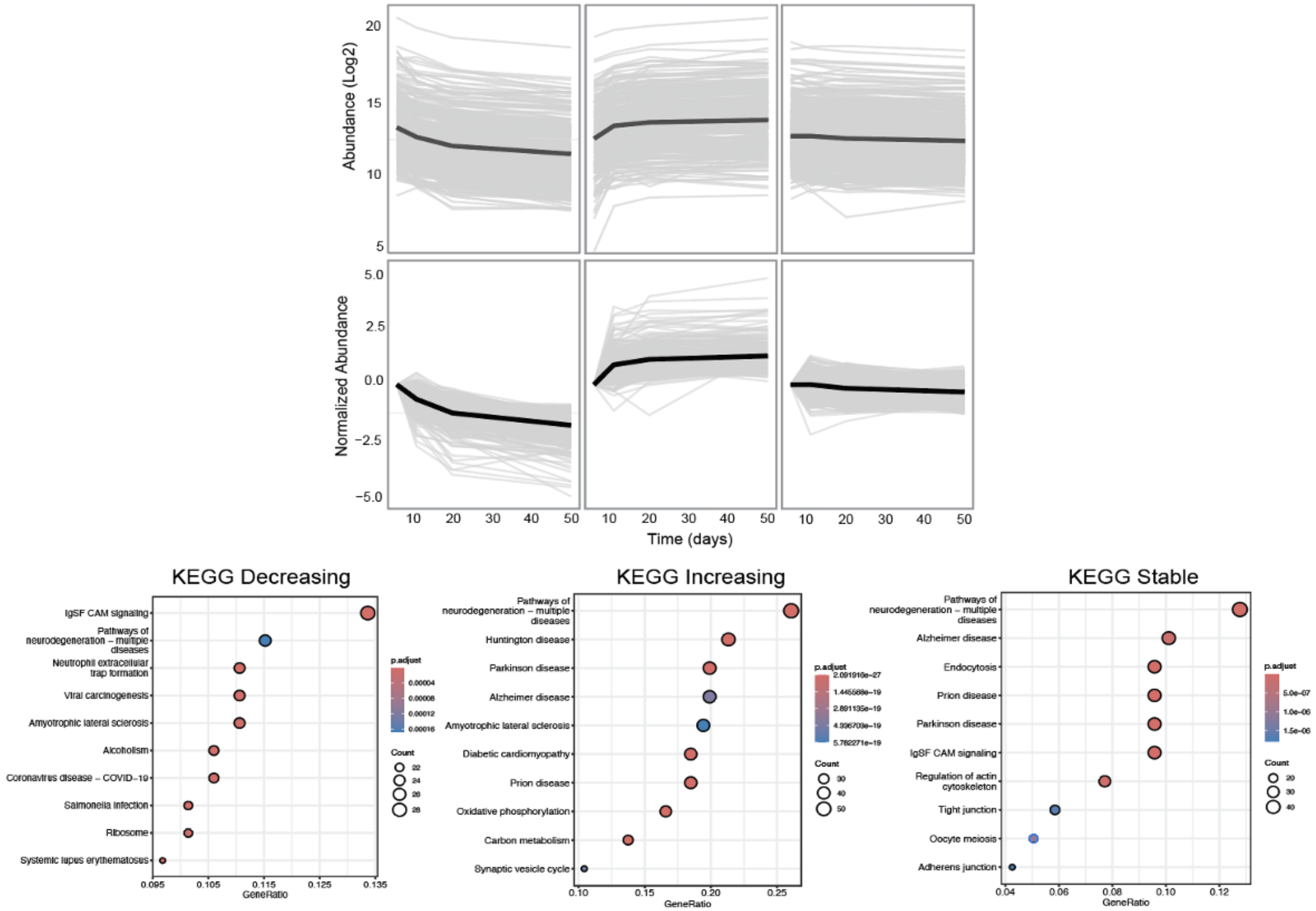


**Supplemental Figure 3: Individual trajectories for proteins across groups with KEGG pathway enrichment**

A) *Top:* Log2 Abundance values for each protein separated by group assignment. Average abundance for group at each timepoint depicted by the black trendline. *Bottom:* same as *top,* but normalized to the P6 timepoint.

B) Top 10 results for each trajectory group by GeneRatio for KEGG pathway overrepresentation analysis. Dots are ranked by GeneRatio, the fraction of genes found in this data set compared to total genes in a term. Dot size and color reflect number of genes and FDR.

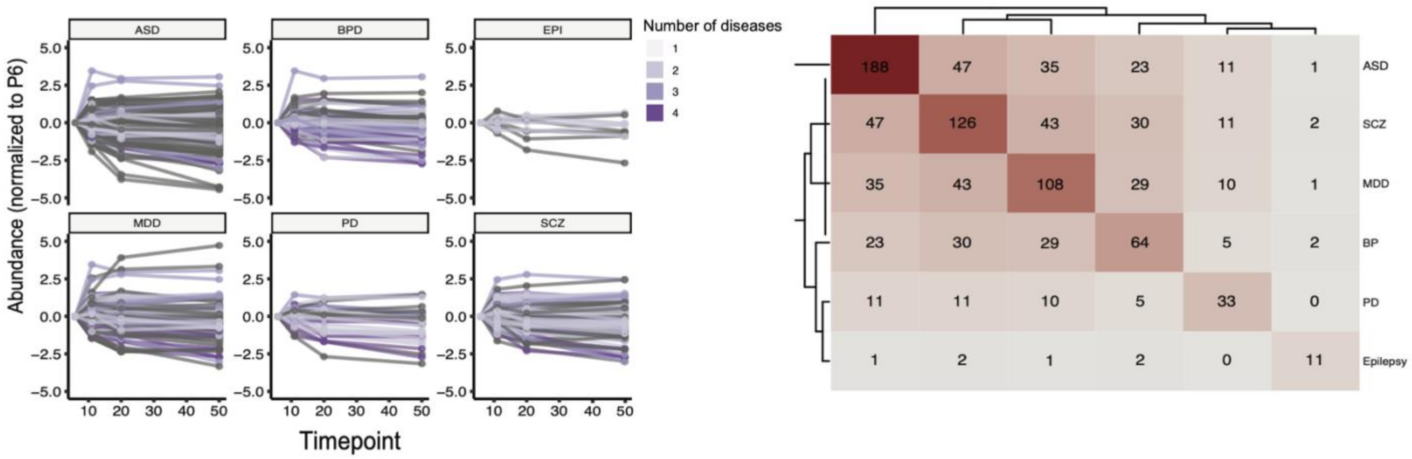


**Supplemental Figure 4: Disease risk allele overlap by count**

A) Log2 abundance values for each disease risk protein, color coded for number of diseases each protein is a risk allele for. Darker purple represents higher degree of overlap.

B) Heatmap representation of overlap counts between risk alleles identified in dopamine axon data. Cells are labeled with the number of shared risk proteins; darker red corresponds to more overlap.
